# APOE4 Accelerates Menopause-Associated Brain Metabolic Shift and Disrupts Bioenergetic Adaptation

**DOI:** 10.64898/2026.03.11.710133

**Authors:** Tian Wang, Yuan Shang, John W. McLean, Fei Yin, Roberta Diaz Brinton

**Author notes:** These authors contributed equally to this work (Tian Wang, Yuan Shang).

## Abstract

**Introduction:** Disruption of brain glucose and lipid metabolism contributes to Alzheimer’s disease (AD) and often emerges before clinical symptoms. Women are at elevated AD risk due to menopause-associated estrogen decline, which impairs mitochondrial function and glucose metabolism. Women’s risk of AD is further elevated by the APOE4 allele, the strongest genetic risk factor for late-onset AD.

**Methods:** To investigate the impact of *APOE* genotype on the menopausal metabolic transition, brain metabolomic and lipidomic profiling was conducted in humanized female APOE3/3, APOE3/4, and APOE4/4 mice across chronological and endocrinological stages of peri-to postmenopausal transition.

**Results:** APOE3/3 mice exhibited dynamic regulation of brain metabolic systems that supported postmenopausal bioenergetic demand. In contrast, APOE3/4 and APOE4/4 mice displayed accelerated and altered metabolic shifts, resulting in postmenopausal amino acid depletion, reduced tricarboxylic acid (TCA) cycle intermediates, lipid accumulation, and alterations in brain lipid composition. A single APOE4 allele was sufficient to impair metabolic adaptation, while APOE4 homozygosity resulted in greater severity of deficits.

**Discussion:** Outcomes of these analyses revealed that APOE4 accelerated menopause-related metabolic decline and compromised bioenergetic adaptation, providing a mechanistic basis for increased AD susceptibility and earlier onset in APOE4-positive women.

## 1. Introduction

Female sex, age, and *APOE4* genotype are the greatest risk factors for Alzheimer’s disease (AD) (Altmann et al., 2014, Farrer et al., 1995, Hebert et al., 2013, Sala Frigerio et al., 2019, Payami et al., 1996, Barnes et al., 2005, Nebel et al., 2018). Women account for two-thirds of AD cases, with increased vulnerability linked to menopausal transitions (2024 Alzheimer’s disease facts and figures, 2024, Brinton, 2008, Brookmeyer et al., 2007, Morrison et al., 2006, Brookmeyer et al., 1998, Brinton et al., 2015). During this period, decline in brain glucose metabolism and mitochondrial function, white matter lipid catabolism, and rise in neuroinflammation contribute to increased AD risk (Yin et al., 2015, Wang et al., 2020, Brinton et al., 2015, Klosinski et al., 2015, Bacon et al., 2019, Mosconi et al., 2017a, Mosconi et al., 2017b). This vulnerability is further amplified in APOE4 carriers, particularly women, who experience earlier symptom onset and accelerated cognitive decline relative to men (Mishra and Brinton, 2018, Mishra et al., 2022, Shang et al., 2020, Wang and Brinton, 2016, Riedel et al., 2016, Farrer et al., 1997, Neu et al., 2017, Belloy et al., 2019, Polsinelli et al., 2023, Holland et al., 2013).

Our recent analyses revealed that APOE4 was associated with earlier age of menopause in women, and that APOE4 women experiencing early menopause exhibited the highest risk of Alzheimer’s (Wang et al., 2025). Mechanistically, using a humanized APOE perimenopause (PAM) mouse model that met human STRAW criteria and was also consistent with human female brain imaging outcomes (Soules et al., 2001, Yin et al., 2015, Wang et al., 2020, Mishra et al., 2020, Mosconi et al., 2021, Mosconi et al., 2017a, Mosconi et al., 2018, Bacon et al., 2019, Brinton et al., 2015, Mishra and Brinton, 2018), we demonstrated that APOE4 carriers failed to mount metabolic reprogramming during the menopausal transition to sustain bioenergetic demands of the brain (Wang et al., 2025). These results are consistent with previous reports that APOE4 impairs brain insulin signaling, disrupts glucose metabolism, and impairs lipid homeostasis and transport (Wu et al., 2018, Qi et al., 2021, Zhao et al., 2017).

Maintaining metabolic homeostasis is essential for brain functions such as synaptic activity, plasticity, energy balance, and neuroprotection (Cunnane et al., 2020). In AD, though a multifactorial disorder, metabolic dysregulation, particularly in glucose and lipid metabolism, plays a crucial role in its pathogenesis (Yin, 2023, Butterfield and Halliwell, 2019). Evidence indicates that disruptions in brain energy metabolism, mitochondrial function, and lipid balance emerge before clinical symptoms and may contribute to the prodromal phase of AD (Yan et al., 2020, Yao et al., 2009, Yin, 2023). Importantly, these metabolic disturbances are significantly influenced by both sex and genetics (Ferretti et al., 2018, Lopez-Lee et al., 2024, Raulin et al., 2022). Notably, APOE4 females display plasma signatures enriched in altered phosphatidylcholines (PCs), suggesting a distinct metabolic phenotype (Chang et al., 2023, Arnold et al., 2020).

Building on these findings, our observed failure of metabolic reprogramming in APOE4 female brains during menopause (Wang et al., 2025) led us to investigate the impact of APOE4 on brain metabolic transitions across the stages of menopause. To elucidate the mechanistic pathways underlying the increased AD risk in APOE4 postmenopausal females, we conducted a comprehensive analysis of global metabolomic and lipidomic profiles in brains from the humanized APOE PAM model. Outcomes indicated that APOE3/3 mice exhibited dynamic regulation of brain metabolic profiles to sustain postmenopausal bioenergetic demand, consistent with previous findings in wild-type mouse and rat PAM models (Wang et al., 2020, Klosinski et al., 2015, Yin et al., 2015). In contrast, APOE3/4 and APOE4/4 mice exhibited accelerated and altered menopause-associated metabolic reprogramming compared to APOE3/3, most notably significant disruptions in lipid metabolism. These shifts resulted in postmenopausal amino acid depletion, reduced TCA cycle metabolites, lipid accumulation, and changes in brain lipid composition. Notably, even a single APOE4 allele was sufficient to confer risk, while APOE4/4 mice displayed the most pronounced dysregulation. Collectively, these findings provide mechanistic insight into increased AD vulnerability and earlier disease onset observed in APOE4-positive women.

## 2. Materials and Methods

### 2.1 Animals (Perimenopausal Animal Model)

All animal studies were performed following National Institutes of Health guidelines on the use of laboratory animals, and all protocols were approved by the University of Arizona Institutional Animal Care and Use Committee. Humanized APOE4/4 targeted replacement (APOE4/4) homozygous mice were obtained from Jackson Laboratory (#027894). Humanized APOE3/wt targeted replacement heterozygous mice were obtained from Jackson Laboratory (#029018) and bred to get homozygous APOE3/3 mice. Humanized APOE3/4 mice were generated by crossing APOE4/4 mice with APOE3/3 mice. Mice were housed on 14 h light/ 10 h dark cycles and provided *ad libitum* access to food (NIH7913) and water.

The estrous cycle status of 6-, 9- and 15-month-old APOE3/3, APOE3/4 and APOE4/4 female mice were monitored by daily vaginal cytology continuously for 3 weeks. Vaginal smears were obtained between 0800 and 1100 h. Four stages of estrous cycle: Estrus (E), Metestrus (M), Diestrus (D) and Proestrus (P), were morphologically characterized based on the proportion of different cell types present in the smears as previously described (Wang et al., 2020, Yin et al., 2015, Wang et al., 2025). Female middle-aged mice were then stratified into 3 different endocrine aging groups with defined stages as per STRAW criteria (Harlow et al., 2012): regular cyclers (4-5 day cycles), irregular cyclers (6-9 day cycles), and acyclic (no cycling within 9 days). Typically, mice transit from regular cycles to irregular cyclers at around 9 months (Finch, 2014). Thus, to capture this endocrinological transition, separate age-matched cohorts of APOE3/3, APOE3/4, and APOE4/4 female mice were monitored, characterized, and collected at 3 time points: 6 months (young, 6M-Reg), 9 months (early perimenopausal transition, 9M-Reg and 9M-Irreg) and 15 months (late perimenopausal transition, 15M-Irreg and 15M-Acyc) for this study (Wang et al., 2025). Mice that did not meet the endocrine status criteria were excluded from the study.

### 2.2 Brain tissue collection

Mice were subjected to an overnight fast lasting approximately 12-16 hours, during which only water was provided *ad libitum*. Anesthesia was administered using inhaled isoflurane, after which the animals were transcardially perfused with cold phosphate-buffered saline (PBS) for 10 minutes. Brains were removed and dissected on ice, snap-frozen on dry ice, and stored in -80°C for subsequent assays.

### 2.3 Metabolomic analysis

Metabolomic analysis on cortex samples was performed by Metabolon utilizing their Global and Complex Lipids Platforms (Wang et al., 2020) (n=5-6/group). Following imputation of missing values, with the minimum observed value for each compound, and log transformation, Welch’s two-sample *t*-tests were used to identify biochemicals that differed significantly between experimental groups within each matrix. Fold change was calculated as the ratio of the mean scaled intensity for a metabolite between two experimental groups.

### 2.4 Heatmap

Heatmaps were generated from group-averaged log-transformed concentrations. Natural log was used for global metabolomics and log2 for lipidomics to better reflect fold changes. Data were then standardized using a Standard Scaler to ensure comparability across metabolites and prevent high-abundance features from dominating the color scale.

### 2.5 Pathway Enrichment

Metabolite Set Enrichment Analysis (MSEA) was performed to determine whether specific metabolic pathways were significantly altered. Log-transformed data were used as input to identify differentially expressed metabolites between groups using the limma (v3.62.2) (Ritchie et al., 2015) R package. The ranked list generated by limma was then analyzed with fgsea (v1.32.4) (Korotkevich et al., 2021) to compute enriched metabolite sets. The fgsea package utilizes a fast, preranked Gene Set Enrichment Analysis algorithm based on a Kolmogorov-Smirnov-like test to determine if a group of metabolites is non-randomly distributed in the ranked list. Normalized enrichment scores (NES) and adjusted p-values (p.adj) were reported.

### 2.6 Correlation analysis

To assess relationships between individual metabolites, correlation analysis was performed on the log-transformed data.

To analyze correlations between metabolite clusters, we summarized the abundance trends within each cluster using an eigenmetabolite (Alter et al., 2000), defined as the first principal component (PC1) of the cluster’s intensity matrix. For each cluster, we constructed a matrix containing the scaled expression values of its metabolites across all samples. Principal Component Analysis (PCA) was then performed on this matrix using the singular value decomposition (SVD) method. The first principal component, which captures the greatest variance within the cluster, was designated as the eigenmetabolite vector.

To ensure that the eigenmetabolite accurately reflected the overall abundance trend of the cluster, we calculated the average intensity profile of the cluster’s metabolites for each sample and computed the Pearson correlation between this mean profile and the PC1 vector. If the correlation was negative, we flipped the sign of the eigenmetabolite vector to align it with the dominant direction of metabolite abundance. All analyses were conducted in Python (v3.10) using the scikit-learn library.

Pearson correlation coefficients were then calculated for all pairwise metabolite comparisons to quantify the strength and direction of linear associations. Statistical significance was assessed using two-tailed t-tests, and correlations with p-values < 0.05 were considered significant.

### 2.7 Statistical Analysis

Statistical analysis of the metabolomics data was performed following previously published methods (Wang et al., 2020). Welch’s two-sample *t*-tests were used to identify biochemicals that differed significantly between experimental groups within each matrix. Pairwise comparisons were performed between *APOE* genotypes within the same chronological and endocrinological groups, as well as between consecutive chronological and endocrinological stages within each genotype. p-values and q-values were reported (Table S1). In the pathway enrichment analysis, pathways with an adjusted p-value < 0.1 were reported, while those with an adjusted p-value < 0.05 were considered statistically significant.

## 3. Results

### 3.1 APOE4 Influences Brain Metabolic Trajectories Across Chronological and Endocrinological Aging

To investigate the impact of *APOE* genotype on the menopause-associated changes in brain metabolic profile, female APOE3/3, APOE3/4, and APOE4/4 mice were assessed at 6, 9, and 15 months of age. Mice were stratified into five chronological and endocrinological aging groups: 6M-Regular (6M-Reg), 9M-Regular (9M-Reg), 9M-Irregular (9M-Irreg), 15M-Irregular (15M-Irreg), and 15M-Acyclic (15M-Acyc), as previously reported (Wang et al., 2025). Brain metabolic profiles were characterized using the Metabolon Global Platform.

As summarized in Table 1, global metabolite profiling demonstrated that female APOE3/3 brain underwent dynamic metabolic changes across the menopausal transition, with the most pronounced alterations evident in 15M-Irreg and 15M-Acyc groups. At the metabolite level, a greater number of statistically significant metabolites were detected in the 15M-Irreg and 15M-Acyc APOE3/3 groups compared with the 9M-Irreg and 15M-Irreg groups, respectively. Correspondingly, at the pathway level, more extensive metabolic pathway alterations were observed in the 15-month groups (Fig. 1).

**Figure 1.**
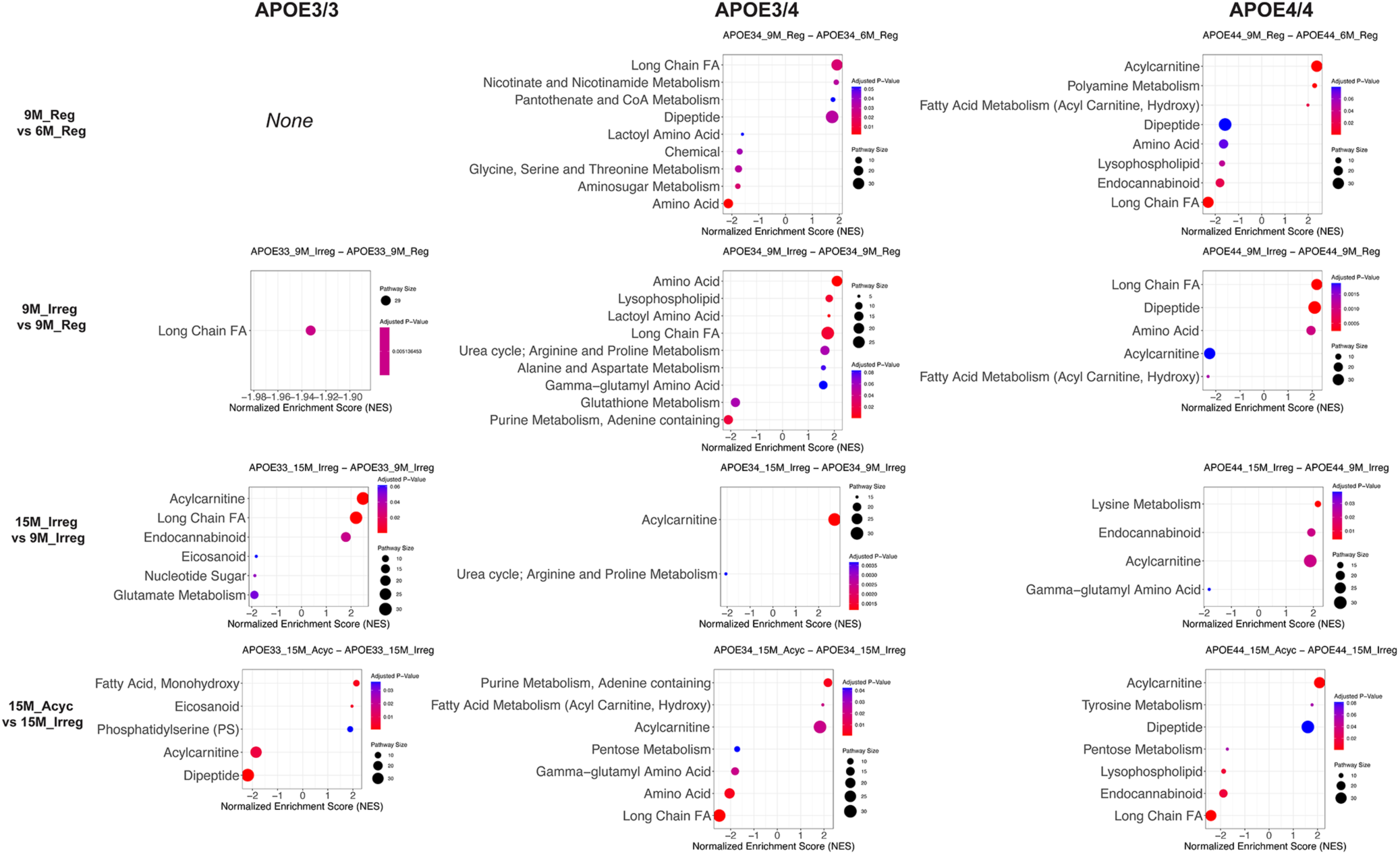
Pathway enrichment analysis comparing chronological and endocrinological groups within each *APOE* genotype. Top enriched metabolic pathways (adjusted p < 0.1) are shown for each pairwise comparison between chronological and endocrinological groups within APOE3/3, APOE3/4, and APOE4/4 groups. Amino acid pathway included the amino acids shown in Figure 3. Long chain fatty acids (FA) pathway included long-chain saturated, monounsaturated, and polyunsaturated fatty acids shown in Figure 4. Acylcarnitine pathway included medium-chain, long-chain saturated, monounsaturated, and polyunsaturated acylcarnitines as shown in Figure 4.

**Table 1.** Summary of the numbers of statistically significant different metabolites (Welch’s two-sample t-test) between chronological and endocrinological groups within each *APOE* genotype (Global Platform).

| Statistical Comparison - Brain (Global) |  |  |  |  |
| --- | --- | --- | --- | --- |
|  | 9M-Reg<br>vs 6M-Reg | 9M-Irreg<br>vs 9M-Reg | 15M-Irreg<br>vs 9M-Irreg | 15M-Acyc<br>vs 15M-Irreg |
| APOE3/3 | 48 | 12 | 89 | 68 |
| APOE3/4 | 37 | 82 | 20 | 16 |
| APOE4/4 | 16 | 48 | 35 | 15 |

In contrast, APOE4 carriers (APOE3/4 and APOE4/4 females) exhibited accelerated metabolic shifts, with more substantial changes observed in the 9M-Irreg group compared with their 9M-Reg counterparts at the metabolite level (Table 1), accompanied by greater pathway-level alterations across the 9-month groups (Fig. 1). These outcomes replicated our findings in both human and mice and were consistent with accelerated endocrine aging in APOE4/4 females compared to APOE3/3 females (Wang et al., 2025).

Further, at the pathway level, although APOE3/4 and APOE4/4 profiles were more similar to each other than to APOE3/3, each genotype exhibited distinct patterns of metabolic alterations (Fig. 1 & 2). When comparing across genotypes, the most pronounced differences were detected in the 15M-Acyc groups between APOE3/3 and APOE3/4 or APOE4/4 and in the 6M-Reg groups between APOE3/4 and APOE4/4 (Fig. 2).

**Figure 2.**
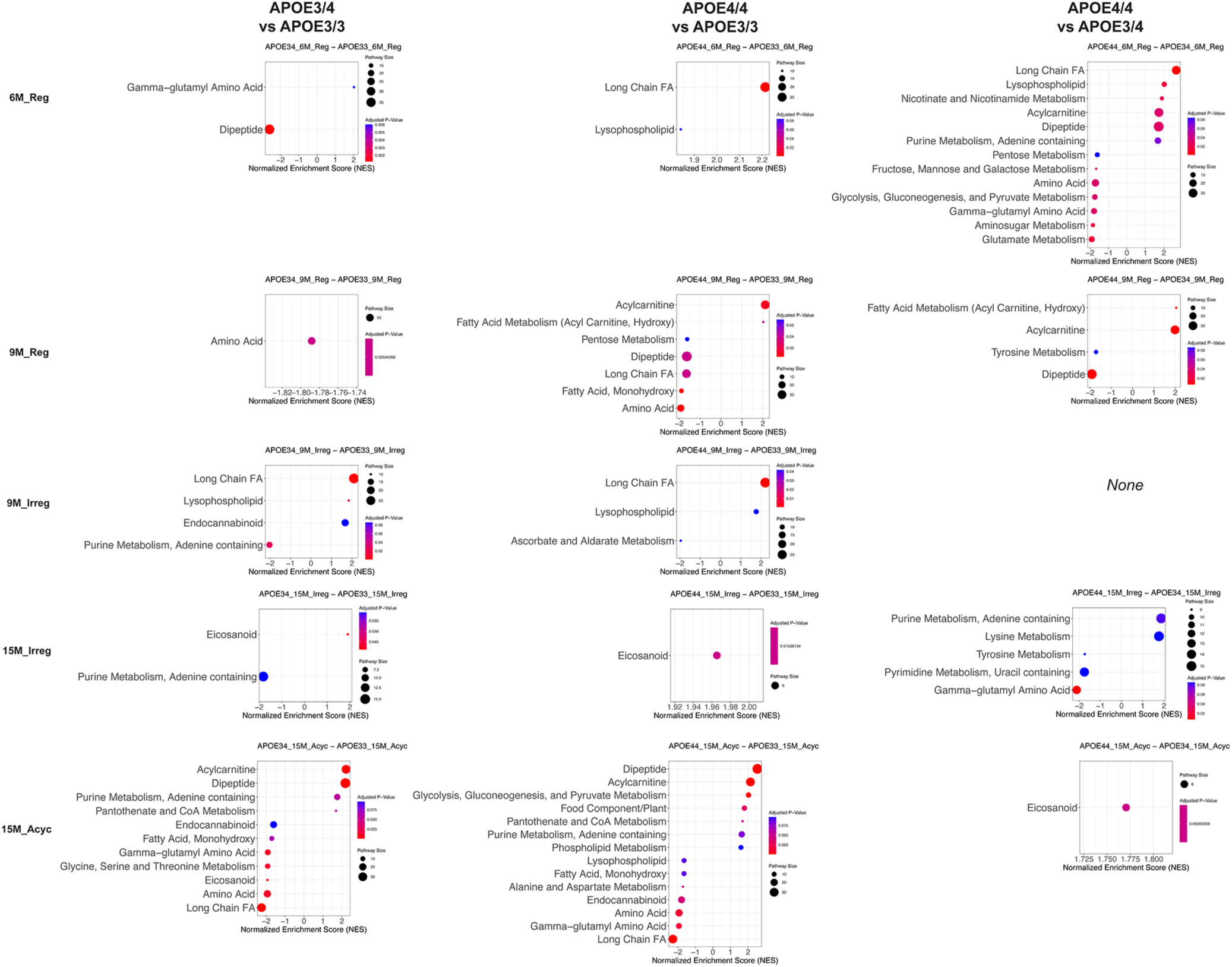
Pathway enrichment analysis of *APO*E genotypes across chronological and endocrinological groups. Top enriched metabolic pathways (adjusted p < 0.1) are shown for each pairwise comparison between *APOE* genotypes across chronological and endocrinological groups. Amino acid pathway included the amino acids shown in Figure 3. Long chain fatty acids (FA) pathway included long-chain saturated, monounsaturated, and polyunsaturated fatty acids shown in Figure 4. Acylcarnitine pathway included medium-chain, long-chain saturated, monounsaturated, and polyunsaturated acylcarnitines as shown in Figure 4.

Taken together, these findings indicated a significant role of the *APOE4* genotype in modulating brain metabolic aging. APOE3/4 and APOE4/4 mice exhibited similar global metabolic aging trajectories, suggesting that even a single APOE4 allele was sufficient to drive these metabolic changes. Furthermore, both APOE3/4 and APOE4/4 mice showed evidence of accelerated metabolic aging compared to APOE3/3 mice.

### 3.2 Amino acid metabolism

Across chronological and endocrinological aging, APOE3/3 females exhibited overall stable brain amino acid and gamma-glutamyl amino acid levels (Fig. 3A, 3B). In contrast, significant alterations in amino acid metabolism were observed in APOE3/4 and APOE4/4 mice. In both genotypes, amino acid pathway, as indicated by enrichment scores (Fig. 1) and supported by individual metabolite levels (Fig. 3A, Table S1), was significantly or near-significantly reduced in the 9M-Reg group compared to the 6M-Reg group. This was followed by a significant increase in the 9M-Irreg group, and a subsequent decline in the 15M groups in both APOE3/4 and APOE4/4 mice (Fig. 1, 3A, Table S1). Similarly, gamma-glutamyl amino acid pathway was elevated in 9M-Irreg APOE3/4 and APOE4/4 females and subsequently decreased significantly in the corresponding 15M groups (Fig. 1, 3B, Table S1). As a result, across genotypes, APOE3/4 and APOE4/4 mice exhibited significantly reduced amino acid pathway in the 9M-Reg groups, which were reversed in the 9M-Irreg groups, followed by a subsequent significant decline accompanied by reduced gamma-glutamyl amino acid pathway in the 15M-Acyc groups compared to their APOE3/3 counterparts (Fig. 2, 3A, 3B, Table S1). These findings indicate that a single or double copy of the APOE4 allele was associated with a pronounced menopause-related upregulation of amino acid metabolism during the early menopausal transition, followed by postmenopausal depletion.

**Figure 3.**
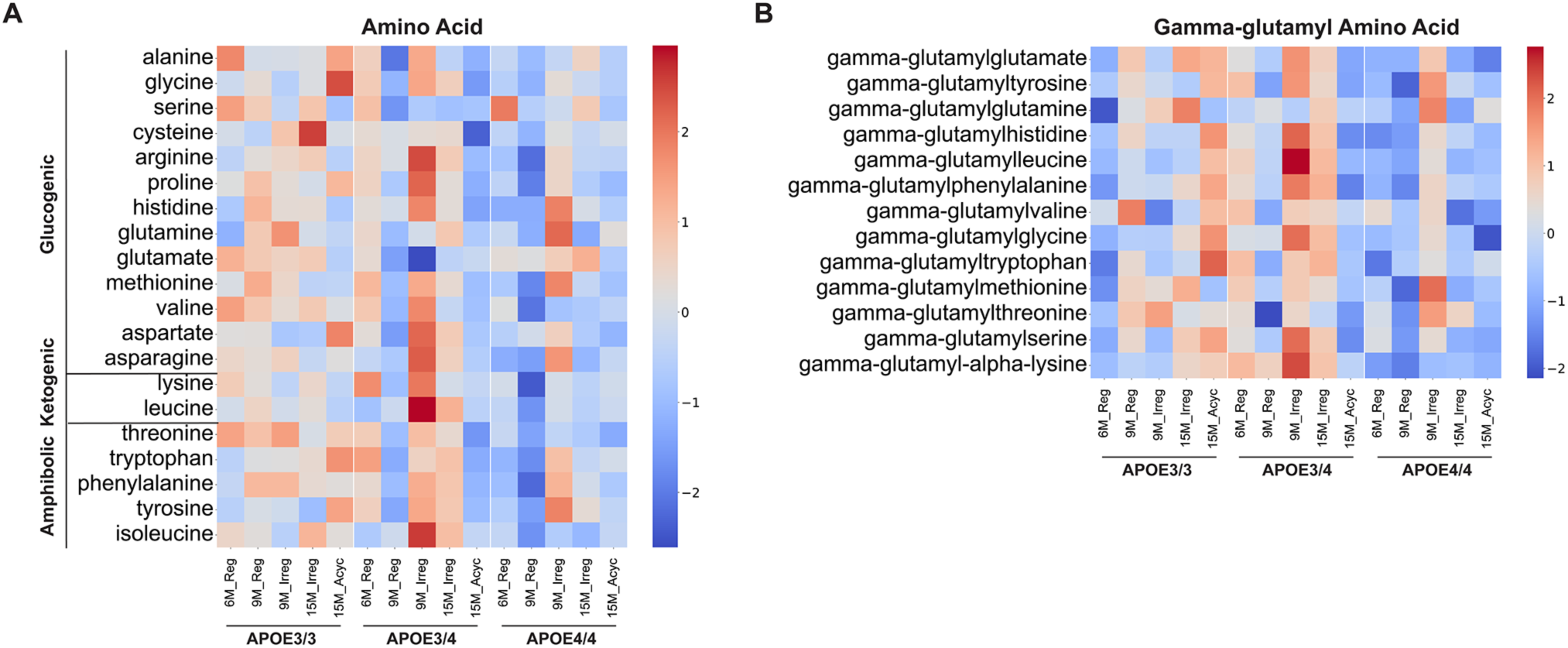
Amino acid metabolism. A. Heatmap of brain amino acid levels. B. Heatmap of brain gamma-glutamyl amino acid levels.

### 3.3 Glycolysis and fatty acid metabolism

The perimenopausal transition is associated with reduced estrogenic control of glucose metabolism and a compensatory upregulation of lipid metabolic pathways in brain, which together trigger downstream cascades that contribute to the activation of white matter catabolism and neuroinflammation (Ding et al., 2013a, Ding et al., 2013b, Mosconi et al., 2017b, Rettberg et al., 2016, Yao et al., 2009, Yin et al., 2015, Klosinski et al., 2015, Mishra et al., 2020, Wang et al., 2020, Mosconi et al., 2017a). Consistently, APOE3/3 15M-Acyc group exhibited reduced pyruvate levels (p=0.047, Table S1) compared to the 15M-Irreg group, with 1,5-anhydroglucitol (1,5-AG), glucose, glucose 6-phosphate, and dihydroxyacetone phosphate (DHAP) also showing decreased abundance (fold change≤0.8, Fig. 4A, Table S1). In contrast, long-chain fatty acid (LCFA) and acylcarnitine pathways were significantly elevated in the 15M-Irreg APOE3/3 group, followed by a significant decline in acylcarnitine pathway in the 15M-Acyc group (Fig. 1, 4B, 4C, Table S1). These findings suggest that, in the APOE3/3 brain, reduced glucose metabolism is accompanied by increased fatty acid oxidation, likely serving as an alternative energy source during the menopausal transition (Klosinski et al., 2015).

**Figure 4.**
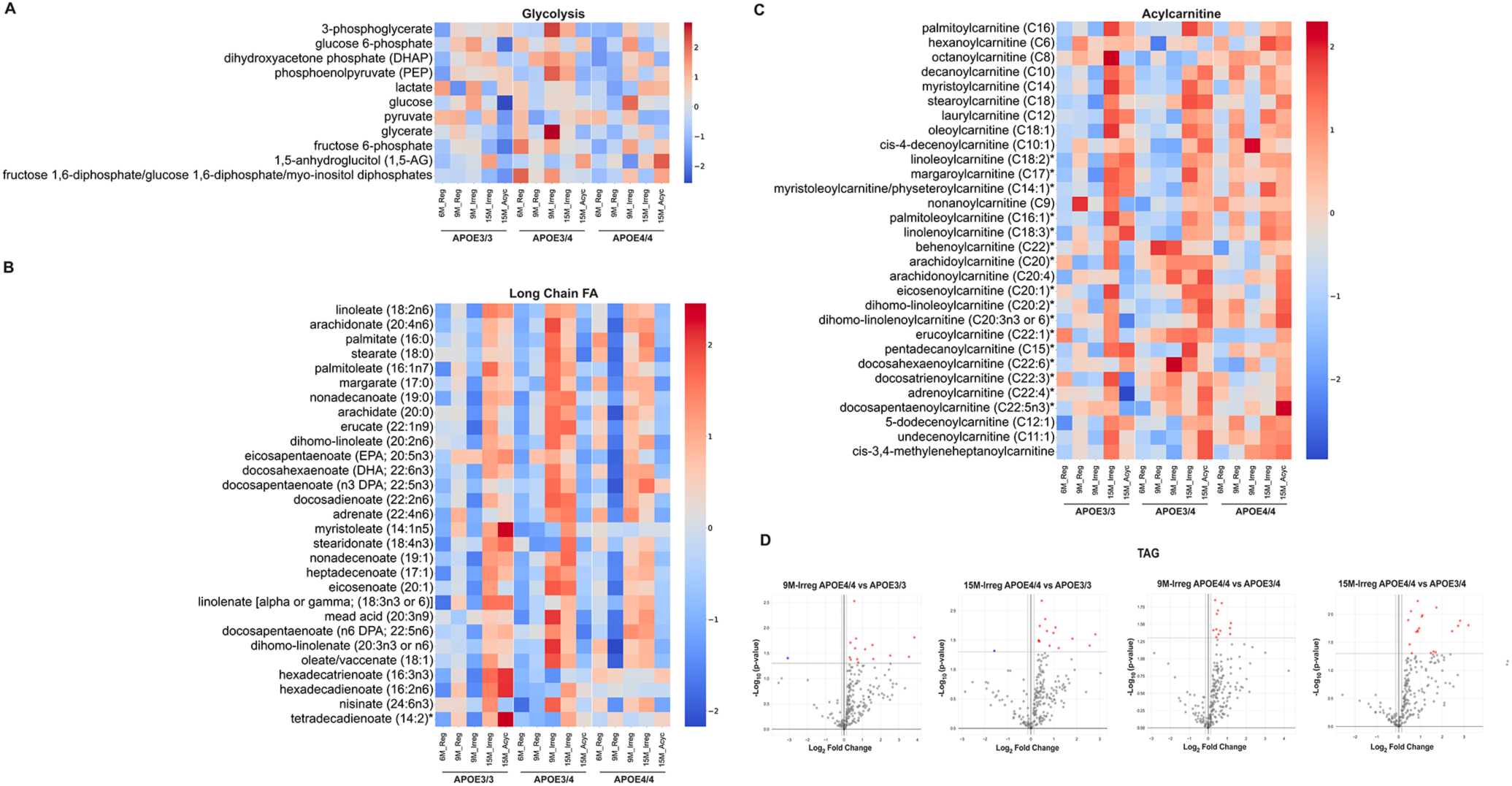
Glucose and lipid metabolism. A. Heatmap of glucose and glycolytic intermediate levels. B. Heatmap of long-chain fatty acid levels. C. Heatmap of acylcarnitine levels. Metabolite name*: Indicated a compound that had not been confirmed based on a standard, but Metabolon was confident in its identity. D. Volcano plots of triacylglycerols (TAGs) comparing the following groups: 9M-Irreg APOE4/4 vs APOE3/3, 9M-Irreg APOE4/4 vs APOE3/4, 15M-Irreg APOE4/4 vs APOE3/3, and 15M-Irreg APOE4/4 vs APOE3/4.

Notably, the presence of the APOE4 allele significantly altered the chronological and endocrinological trajectories of brain metabolites involved in glycolysis and fatty acid metabolism. Specifically, glycolysis-related metabolites remained stable in the 15M APOE3/4 and APOE4/4 groups, resulting in significant elevation of the glycolysis metabolism pathway in 15M-Acyc APOE4/4 brains compared to their APOE3/3 counterparts (Fig. 2, 4A).

In parallel, changes in LCFA profiles were highly dynamic across the menopausal transition, irrespective of *APOE* genotype. However, the key distinction among *APOE* genotypes was the timing of these shifts. In APOE3/3 females, LCFA levels increased later in the menopausal process. In contrast, both APOE3/4 and APOE4/4 females exhibited LCFA profiles consistent with earlier utilization during the endocrine aging transition. Specifically, LCFA pathway was significantly elevated in 6M-Reg APOE4/4 mice compared to APOE3/3 6M-Reg group. In APOE4/4 mice, LCFA pathway declined at 9M-Reg relative to 6M-Reg, followed by a significant rebound in the 9M-Irreg group, which remained elevated in the 15M-Irreg group, and subsequently declined significantly in the 15M-Acyc group (Fig. 1, 4B, Table S1). In parallel, acylcarnitine pathway in APOE4/4 mice significantly increased at 9M-Reg, decreased at 9M-Irreg, and rebounded at 15 months (Fig. 1, 4C, Table S1). A similar pattern also occurred in APOE3/4 females, with an earlier increase in LCFAs (9M) and delayed elevation in acylcarnitines (15M) (Fig. 1, 4B, 4C, Table S1). This temporal mismatch between LCFA accumulation and acylcarnitine response was indicative of dysregulated fatty acid metabolism and a diminished capacity for menopause-induced lipid adaptation in APOE4 carriers.

Further, at the 15M-Acyc stage, both APOE3/4 and APOE4/4 females exhibited significantly reduced LCFA pathway and elevated acylcarnitine pathway compared to APOE3/3 counterparts (Fig. 2, 4B, 4C, Table S1). Specific to the APOE4/4 brain was the marked upregulation of triacylglycerol (TAG) metabolites at the 9M-Irreg and 15M-Irreg stages relative to APOE3/3 and APOE3/4 counterparts (Fig. 4D).

These findings are consistent with preserved glycolytic metabolism but disrupted lipid metabolism regulation in APOE4 carriers, with more pronounced impairment in APOE4/4 brains, leading to increased lipid accumulation in the postmenopausal APOE4/4 brain.

### 3.4 Energy metabolism

The brain relies on ATP primarily generated in mitochondria through oxidative phosphorylation of glucose via the tricarboxylic acid (TCA) cycle (Dienel, 2019, Cunnane et al., 2020). In APOE3/3 females, a trend towards increased TCA metabolites was observed in the 15M-Acyc group compared to the 15M-Irreg group (Fig. 5A, Table S1). Specifically, citrate levels were elevated (p=0.041, Table S1), consistent with increases in aconitate, isocitrate, alpha-ketoglutarate, and fumarate (fold change ≥1.2) in APOE3/3 15M-Acyc brains relative to APOE3/3 15M-Irreg brains (Fig. 5A, Table S1). These findings are consistent with the activation of TCA-related gene expression previously reported in 15M-Acyc APOE3/3 brains (Wang et al., 2025), indicating menopause-induced mitochondrial reprogramming.

**Figure 5.**
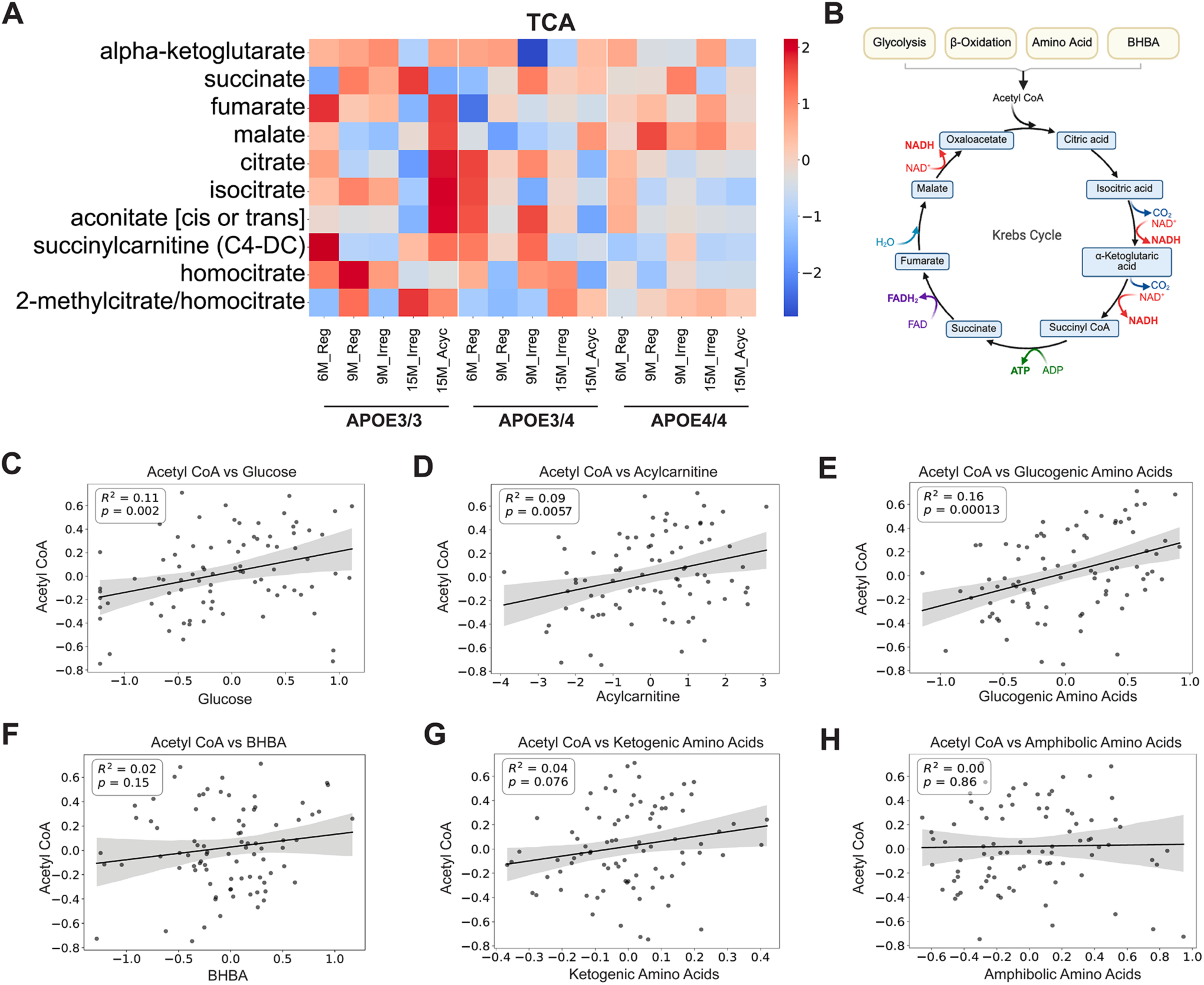
Energy production. A. Heatmap of TCA metabolite levels. B. Pathway diagram (created with BioRender.com). Correlations between acetyl-CoA and key metabolic substrates across groups: (C) glucose, (D) acylcarnitines, (E) glucogenic amino acids, (F) β-hydroxybutyrate (BHBA), (G) ketogenic amino acids, and (H) amphibolic amino acids. The x- and y-axes represent log-normalized metabolite concentrations or metabolite cluster eigenvalues.

In contrast, the postmenopause-associated increase in TCA cycle metabolites was not detected in APOE3/4 or APOE4/4 15M-Acyc groups. Notably, postmenopausal APOE3/4 and APOE4/4 brains exhibited reduced TCA cycle metabolites compared to APOE3/3 brains. Both 15M-Acyc APOE3/4 and APOE4/4 brains exhibited a trend toward reduced levels of citrate, aconitate, isocitrate, alpha-ketoglutarate, and fumarate (fold change ≤0.8) relative to APOE3/3 (Fig. 5A, Table S1), consistent with our previous findings of impaired mitochondrial reprogramming in 15M-Acyc APOE4/4 brains (Wang et al., 2025).

Further, acetyl-CoA functions as the critical entry point into the TCA cycle and can be derived from multiple metabolic pathways, including glycolysis (glucose), β-oxidation of fatty acids, amino acid catabolism, and ketone body metabolism (Neupane et al., 2019) (Fig. 5B). Across all groups, brain acetyl-CoA levels were significantly positively correlated with brain glucose, acylcarnitines, and glucogenic amino acids, but not with ketone bodies (BHBA), ketogenic amino acids, or amphibolic amino acids (Wang et al., 2020) (Fig. 5C-5H). These correlations indicated that the metabolic capacity for glycolysis, β-oxidation, and amino acid catabolism was preserved across chronological and endocrinological aging and *APOE* genotypes, with an overall coupling of glucose, fatty acids, and glucogenic amino acids to the TCA cycle in the brain.

When stratified by *APOE* genotype, distinct patterns of metabolic association were observed (Fig. S1). In the APOE3/3 groups, acetyl-CoA levels showed a significant positive correlation with both glucose and glucogenic amino acids, and a moderate correlation with acylcarnitines, suggesting a broad reliance on multiple fuel sources for acetyl-CoA in these brains. In the APOE3/4 group, acetyl-CoA levels were significantly correlated only with acylcarnitines, suggesting a metabolic shift toward lipid utilization. In contrast, the APOE4/4 group exhibited significant correlations between acetyl-CoA and glucose, glucogenic amino acids, and ketogenic amino acids. Notably, APOE4/4 brains exhibited a marked accumulation of acylcarnitines and TAGs (Fig. 4C, 4D) without a corresponding increase in TCA cycle metabolites, suggesting impaired regulation of β-oxidation, potentially due to either reduced β-oxidation capacity or saturation of the pathway.

Collectively, these results indicate that mitochondrial energy production was reduced in the postmenopausal APOE3/4 and APOE4/4 brain, and that APOE4 further disrupts the integration of diverse fuel sources into the TCA cycle.

### 3.5 Cholesterol metabolism

APOE4 has been reported to cause dysregulation in cholesterol metabolism(Blanchard et al., 2022, Martins et al., 2006); therefore, cholesterol metabolite profiles were analyzed across all groups. An *APOE*-genotype-dependent difference was observed at 9M-Irreg stage (Fig. 6A, Table S1). Specifically, the APOE4/4 9M-Irreg group exhibited elevated levels of cholesterol sulfate, 7alpha-hydroxy-3-oxo-4-cholestenoate (7-Hoca), 4-cholesten-3-one, and 7-hydroxycholesterol (fold change ≥1.2) compared to APOE3/3 counterparts. More moderate changes were observed in the APOE3/4 group, with increased levels of 7-HOCA and 4-cholesten-3-one also observed at 9M-Irreg (Fig. 6A, Table S1). Consistently, cholesterol ester levels were elevated, with total cholesterol ester levels increased 2.70-fold (p=0.042) in 9M-Irreg APOE4/4 group compared to APOE3/3 counterparts (Fig. 6B, Table S1). These findings are consistent with enhanced cholesterol metabolism in the brains of APOE4 carriers during the early perimenopausal transition, particularly in APOE4/4 individuals.

**Figure 6.**
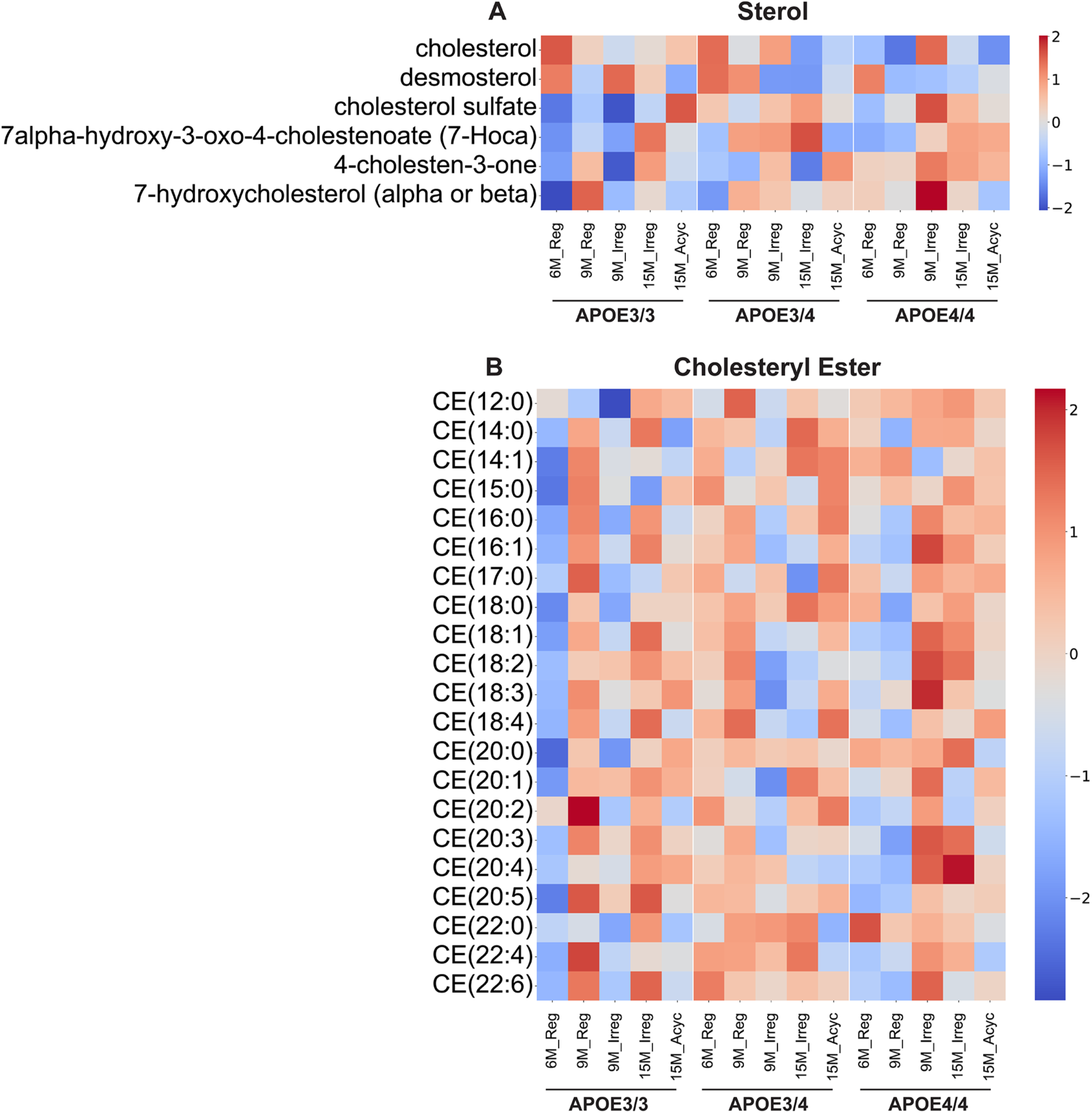
Cholesterol metabolism. A. Heatmap of cholesterol-related metabolite levels. B. Heatmap of cholesterol ester levels.

### 3.6 Brain lipid composition

The major classes of brain lipids include phospholipids, sphingolipids, and neutral complex lipids, each serving distinct yet interrelated roles in maintaining neural homeostasis. Phospholipids and sphingolipids are essential structural components of cellular membranes and are also critical to signal transduction pathways (Farooqui and Farooqui, 2024). In contrast, neutral lipids primarily function as intracellular energy reservoirs and are involved in the storage of fatty acids, particularly under metabolic stress (Yoon et al., 2022, Tracey et al., 2018). To further investigate the impact of *APOE* genotype on brain lipid composition across the menopausal transition, brain lipidomic profiles were characterized using the Metabolon Complex Lipid Platform.

Lipidomic analyses revealed that APOE3/3 females exhibited dynamic shifts in brain lipid species across the menopausal transition, as indicated by a high number of statistically significant lipid species between chronological and endocrinological groups (Table 2). In contrast, APOE3/4 and APOE4/4 brains exhibited fewer significant lipid alterations across both chronological and endocrinological aging. Notably, APOE4 carriers (APOE3/4 and APOE4/4 females) exhibited a shift in lipid composition with a higher proportion of neutral complex lipids, consistent with increased fatty acid content, and a lower proportion of phospholipids in the brain compared to APOE3/3 females (Fig. 7A). Specifically, elevated ceramide levels were observed in the 9M-Irreg APOE3/4 and APOE4/4 groups, with total ceramide levels increased by 1.42- and 1.31-fold, respectively, compared to APOE3/3 counterparts (p=0.024 and 0.045; Fig. 7B, 7C, Table S1). In contrast, a reduction in phosphatidylcholine levels was detected in 15M-Acyc APOE3/4 and APOE4/4 brains, with total levels decreased by 0.85- and 0.92-fold relative to APOE3/3 mice (p=0.295 and 0.038), suggesting compromised membrane integrity (Fig. 7B, 7D, Table S1).

**Figure 7.**
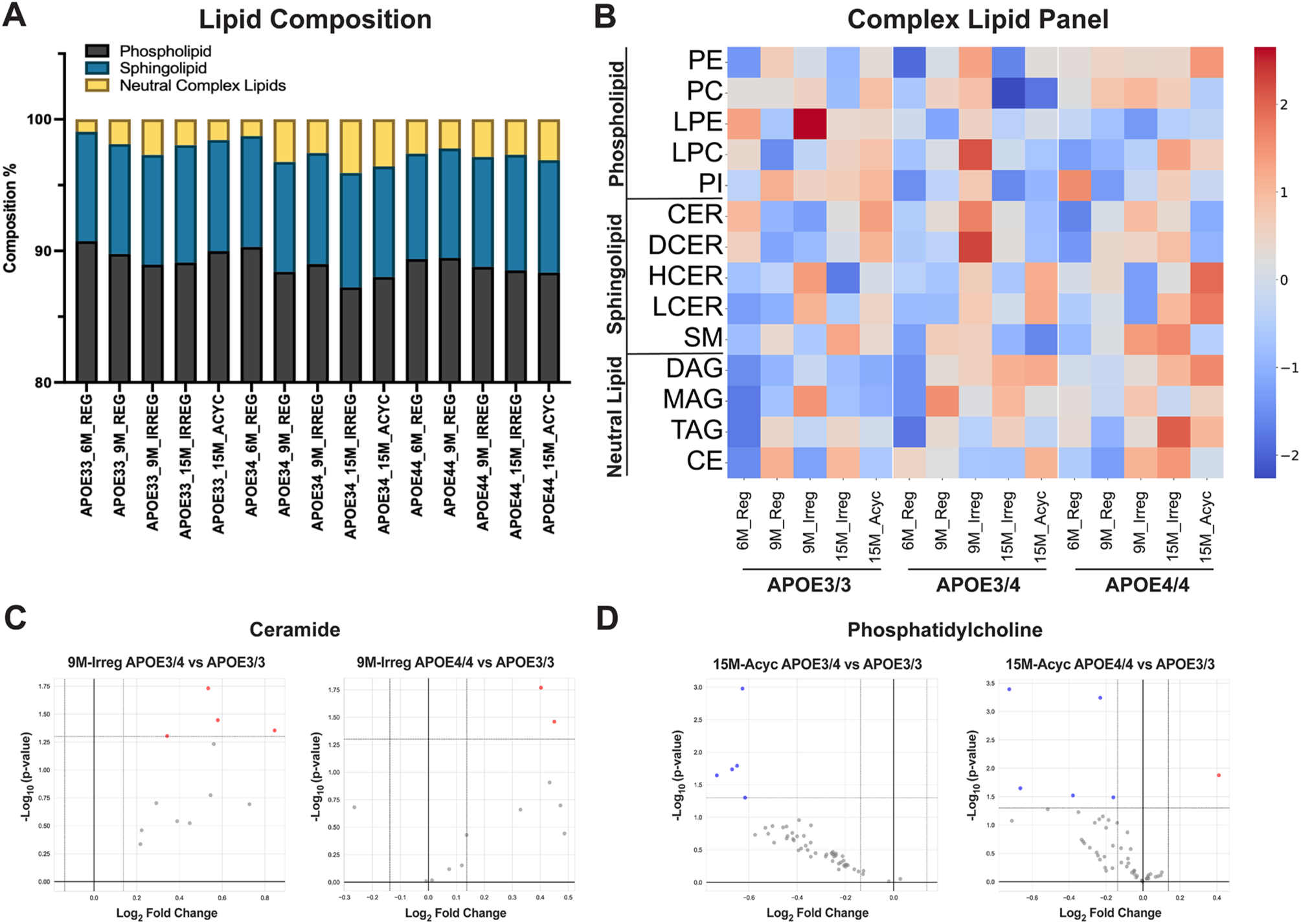
Brain lipidomic profile. A. Brain lipid composition. B. Heatmap of lipid concentrations. C. Volcano plots of ceramides comparing the following groups: 9M-Irreg APOE3/4 vs APOE3/3 and 9M-Irreg APOE4/4 vs APOE3/3. D. Volcano plots of phosphatidylcholines comparing the following groups: 15M-Acyc APOE3/4 vs APOE3/3 and 15M-Acyc APOE4/4 vs APOE3/3.

**Table 2.** Summary of the numbers of statistically significant different metabolites (Welch’s two-sample t-test) between chronological and endocrinological groups within each *APOE* genotype (Complex Lipid Platform).

| Statistical Comparison - Brain (Complex Lipid) |  |  |  |  |
| --- | --- | --- | --- | --- |
|  | 9M-Reg<br>vs 6M-Reg | 9M-Irreg<br>vs 9M-Reg | 15M-Irreg<br>vs 9M-Irreg | 15M-Acyc<br>vs 15M-Irreg |
| APOE3/3 | 59 | 30 | 75 | 28 |
| APOE3/4 | 38 | 7 | 12 | 6 |
| APOE4/4 | 18 | 20 | 16 | 29 |

## 4. Discussion

Both preclinical and human studies indicate that the perimenopausal transition represents a critical tipping point for the onset of a bioenergetic crisis, driven by the decline in estrogenic control of glucose metabolism, impaired mitochondrial function, and a compensatory upregulation of lipid metabolic pathways, leading to increased white matter catabolism and increased neuroinflammation, all of which contribute to increased AD risk in women (Ding et al., 2013a, Ding et al., 2013b, Mosconi et al., 2017b, Rettberg et al., 2016, Yao et al., 2009, Klosinski et al., 2015, Wang et al., 2020, Yin et al., 2015, Mosconi et al., 2017a).

The brain metabolic profiles observed in APOE3/3 PAM models recapitulate the patterns found in wild-type mouse and rat PAM models (Wang et al., 2020, Klosinski et al., 2015, Yin et al., 2015), confirming that the menopause effect is conserved and reproducible across rodent species. In APOE3/3 mice, glucose metabolism decreased postmenopausally, accompanied by increased lipid metabolism and a rebound in TCA cycle metabolites. This profile aligns with the adaptive upregulation of TCA cycle-related gene expression that supports mitochondrial function in response to estrogen loss, along with an increased reliance on lipid metabolism as an alternative energy source, as reported in APOE3/3, wild-type mouse, and rat PAM models (Wang et al., 2020, Yin et al., 2015, Wang et al., 2025, Klosinski et al., 2015). These dynamic metabolic shifts are consistent with the documented role of estrogen in regulating energy metabolism (Rettberg et al., 2014) and highlight the importance of adaptive shifts during menopause in response to estrogen loss to sustain brain function postmenopausally. Furthermore, the APOE3/3 brain exhibited consistent coupling of glucose, glucogenic amino acids and fatty acid metabolism to the TCA cycle across both chronological and endocrinological aging. This suggests that the APOE3/3 brain is able to maintain efficient energy production, consistent with glucose serving as the primary energy source while β-oxidation contributes up to ∼20% of the brain’s energy expenditure (Szrok-Jurga et al., 2023).

In contrast, the distinguishing feature of APOE4 carriers lies in the severity and timing of the metabolic shift, which emerges earlier and with greater intensity. Both APOE3/4 and APOE4/4 mice display exaggerated metabolic changes during the early perimenopausal transition, consistent with accelerated aging (Wang et al., 2025). Importantly, in the postmenopausal groups, APOE3/4 and APOE4/4 brains exhibit metabolic profiles that shift toward an AD-like state (Chang et al., 2023, Batra et al., 2023).

In contrast to APOE3/3 brains, amino acid metabolism increased significantly during early perimenopause in APOE3/4 and APOE4/4 brains, potentially as an alternative energy source (Wang et al., 2020). However, this compensatory pathway is not sustainable, leading to postmenopausal amino acid depletion. In comparison, APOE3/3 brains maintain more stable amino acid concentrations throughout the transition. Amino acids are essential for brain function, acting not only as structural components for protein synthesis but also as neurotransmitters, metabolic substrates, and regulators of redox balance (Ling et al., 2023). The transient compensatory shift toward amino acid oxidation in response to reduced glucose metabolism has been implicated in AD, where subsequent disturbances in amino acid levels and their catabolites may contribute to disease progression (Griffin and Bradshaw, 2017). Consistent with this, the menopause-associated amino acid metabolism alterations observed in APOE3/4 and APOE4/4 brains may contribute to impaired synthetic function and the increased AD risk in postmenopausal APOE4 women.

Further, APOE3/4 and APOE4/4 brains fail to mount the adaptive transcriptional reprogramming, leading to reduced TCA metabolites postmenopausally, consistent with impaired mitochondrial function observed in both animal and human APOE4 studies, thereby contributing to increased AD risk (Area-Gomez et al., 2020, Shang et al., 2020, Hesse et al., 2019, Yin et al., 2020, Wang et al., 2025). Instead, postmenopausal APOE4/4 brains displayed preserved glycolytic capacity and elevated glycolytic metabolites, which may reflect reduced flux into the TCA cycle and a compensatory upregulation of glycolysis to sustain energy production under conditions of mitochondrial dysfunction. This is consistent with evidence of enhanced glycolysis in APOE4 neurons (Orr et al., 2019) and astrocytes (Qi et al., 2021), as well as in young female APOE4 carriers (Farmer et al., 2021), and with findings of glucose accumulation and impaired glycolytic flux in AD brains (An et al., 2018).

The most profoundly disrupted metabolic pathway in APOE4 carriers is lipid metabolism. APOE4 is known to impair cholesterol metabolism, disrupt lipid transport, increase triacylglycerol accumulation, and alter lipid composition and myelination (Qi et al., 2021, Martins et al., 2006, Huang and Mahley, 2014, Blanchard et al., 2022, Shang et al., 2020). Our results demonstrated that APOE3/3 females exhibited a coordinated and dynamic regulation of LCFAs and acylcarnitines throughout the menopausal transition, supporting efficient lipid utilization and metabolic adaptation. In contrast, APOE4/4 brains lacked this adaptive flexibility. APOE4/4 mice exhibited elevated levels of cholesterol metabolites, cholesterol esters, LCFAs, TAGs and ceramides during early perimenopause, followed by increased acylcarnitine levels and sustained TAG elevation in late perimenopause, which persisted into postmenopause, along with a reduction in phosphatidylcholine and LCFA levels. Notably, APOE3/4 brains exhibited similar but milder menopause-driven alterations in lipid profiles and maintained β-oxidation capacity, whereas APOE4/4 brains revealed more pronounced impairments in both β-oxidation and lipid utilization. This disrupted coordination of lipid metabolic pathways is consistent with the impaired lipid transport and β-oxidation capacity of APOE4-expressing astrocytes (Qi et al., 2021), leading to lipid accumulation rather than utilization for energy. Importantly, such disrupted lipid homeostasis not only compromises bioenergetic function and membrane integrity but also promotes neuroinflammation (Haney et al., 2024, Victor et al., 2022, Yang et al., 2023), thereby contributing to AD pathogenesis and modifying disease risk (Yin, 2023). Further, reduced serum phosphatidylcholines have been identified as key metabolic signatures in APOE4-positive AD patients and are strongly associated with cognitive decline (Chang et al., 2023, Arnold et al., 2020).

The compromised metabolic profile elucidated herein provides a mechanistic basis for elevated AD risk in APOE3/4 and APOE4/4 women while also identifying therapeutic targets that could mitigate the impact of the APOE4 genotype. Collectively, these findings provide mechanistic evidence that APOE4 exacerbates the menopause-associated metabolic transition toward a pro-AD phenotype, increasing the vulnerability of APOE4-carrying women to earlier onset and progression of Alzheimer’s disease.

## Data availability statement

The datasets generated and/or analyzed during the current study are available from the corresponding author on reasonable request.

## Acknowledgments

We thank Zisu Mao for their contribution.

---

Figure S1. Correlations between acetyl-CoA and key metabolic substrates across groups in each *APOE* genotype: (A) glucose, (B) acylcarnitines, (C) glucogenic amino acids, and (D) ketogenic amino acids.

