## Supplementary figures and images for "APOE4 Accelerates Menopause-Associated Brain Metabolic Shift and Disrupts Bioenergetic Adaptation"

### Supplemental Figure 1

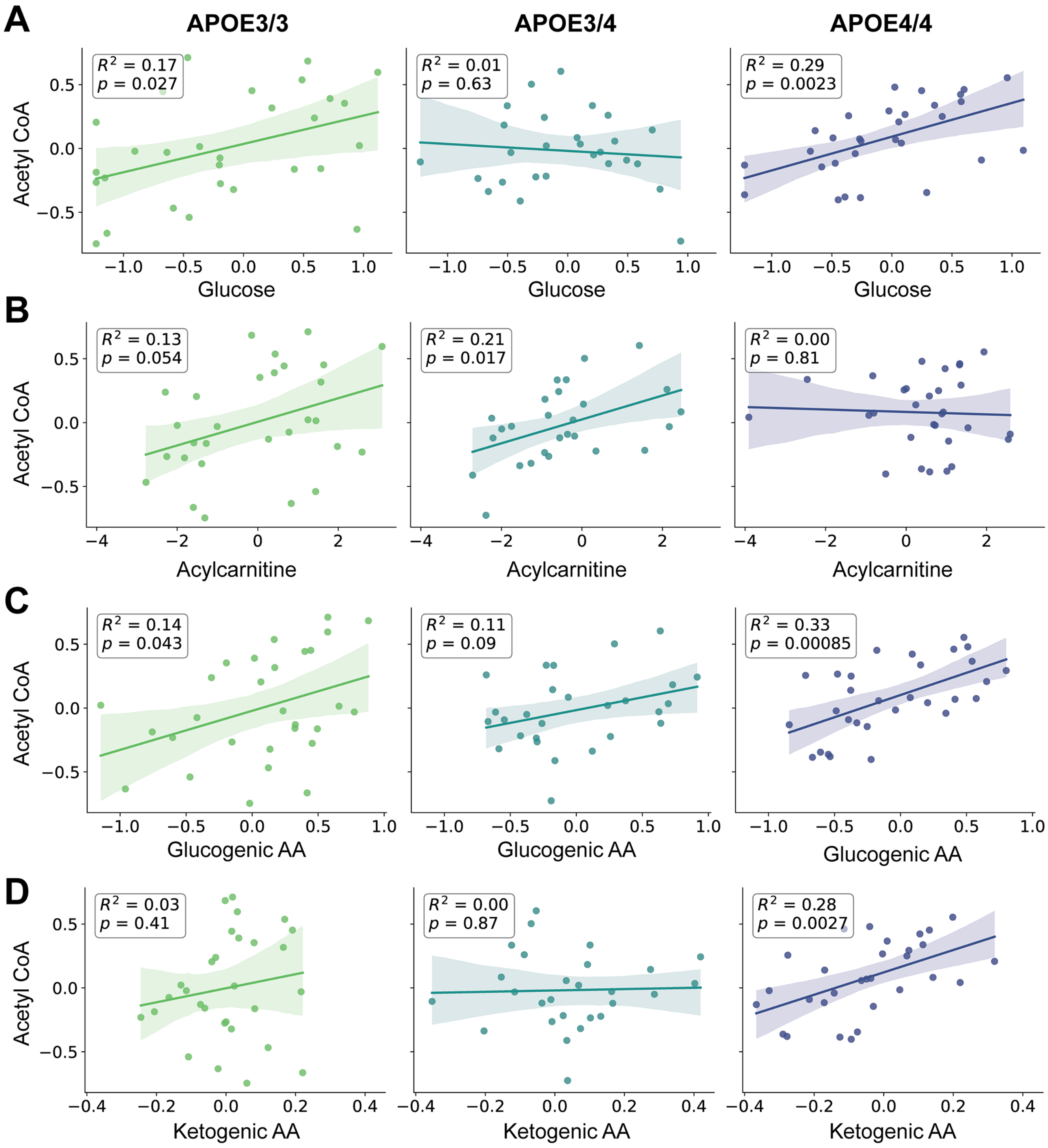
